# Genomic imprinting drives eusociality

**DOI:** 10.1101/2021.04.28.441822

**Authors:** Kenji Matsuura, Hiromu Ito, Kazuya Kobayashi, Haruka Osaki, Jin Yoshimura, Edward L. Vargo

**Affiliations:** Laboratory of Insect Ecology, Graduate School of Agriculture, Kyoto University, Kyoto, Japan.; Department of International Health and Medical Anthropology, Institute of Tropical Medicine, Nagasaki University. Nagasaki, Japan.; Hokkaido Forest Research Station, Field Science Education and Research Center, Kyoto University, Shibecha, Hokkaido, Japan.; Department of Biology, Kyushu University, Fukuoka, Japan.; Department of Entomology, Texas A&M University, College Station, TX, USA.

**Keywords:** origin of eusociality, termite, epigenetic inheritance, sex antagonism

## Abstract

The origin of eusociality, altruistically foregoing personal reproduction to help others, has been a long-standing paradox ever since Darwin. Most eusocial insects and rodents likely evolved from subsocial precursors, in which older offspring “helpers” contribute to the development of younger siblings without a permanent sterile caste. The driving mechanism for the transition from subsociality (with helpers) to eusociality (with lifelong sterile workers) remains an enigma because individuals in subsocial groups are subject to direct natural selection rather than kin selection. Our genomic imprinting theory demonstrates that natural selection generates eusociality in subsocial groups when parental reproductive capacity is linked to a delay in the sexual development of offspring due to sex-antagonistic action of transgenerational epigenetic marks. Focusing on termites, our theory provides the missing evolutionary link to explain the evolution of eusociality from their subsocial wood-feeding cockroach ancestors, and provides a novel framework for understanding the origin of eusociality.

## Introduction

The origin of eusociality poses a major challenge to Darwin’s theory of natural selection (Darwin, 1859) and it remains a subject of debate today (Kay et al., 2020; Nowak et al., 2010; Wilson and Hölldobler, 2005). Eusociality is an advanced state of sociality characterized by the presence of workers that forego personal reproduction in exchange for altruistic behavior toward close relatives. It has long been studied within the paradigm of inclusive fitness and kin selection (Hamilton, 1964). However, the role of kin selection in the origin of eusociality remains controversial (Wilson and Hölldobler, 2005). Explanations of the origin of new traits are faced with the ‘‘selection paradox,’’ the notion that selection cannot act on traits that do not yet exist, and therefore cannot directly cause novelty (Jacob, 1977; Müller and Wagner, 1991). While biologists have made great progress in the last 150 years in understanding how existing traits diversify, we have made less progress in understanding how novel traits arise in the first place. What generates the novelty—here the first appearance of sterile workers?

In colonies of eusocial insect (ants, bees, wasps and termites), workers take on non-reproductive tasks and contribute to their parents’ reproduction (Wilson, 1971). The colony essentially behaves as a superorganism operating as a functionally integrated unit, analogous to the organization of unitary organisms (Hölldobler and Wilson, 2009). In eusocial colonies, the worker caste is an extended phenotype of the parents. Unlike unitary organisms, however, the extended phenotype of a social insect colony enables selection to act at both the individual (within-colony selection) and colony levels (between colony selection). Suppose the first eusocial colony appears on Earth. This colony uniquely has lifelong sterile workers, while all other attributes are identical to other colonies. Initially, the colony cannot compensate for the loss of fitness due to sterility of some offspring unless the parents can simultaneously increase their reproductive output to produce a greater number of fertile offspring. One or more underlying mechanisms are required to link worker sterility with increased reproductive capacity of the parents.

Termite societies, like those of eusocial Hymenoptera (ants and some bees and wasps), are generally large families with a reproductive division of labor. These two groups evolved complex societies independently with different life history pre-adaptations for social evolution. Termites are the oldest social insects, with societies dating back to 150 million years ago (Bourguignon et al., 2015). Eusociality in termites evolved from a subsocial precursor that lived in close family groups and occupied dead trees, as in its sister group Cryptocercidae (woodroaches) (Thorne, 1997) (Fig. 1). The confined nest cavities, slow development due to poor nutritional quality of wood, and need for refaunation of obligate gut symbionts selected for parental care and long-term associations of monogamous family groups (Nalepa, 1994). One requisite condition for the origin of eusociality is iteroparity, which is not present in extant semelparous woodroaches (Korb, 2008; Thorne, 1997). In the subsocial ancestor, iteroparous parents produced brood of staggered age classes living together, where all surviving offspring eventually matured into reproductive imagoes and dispersed to found new colonies.

**Figure 1.**
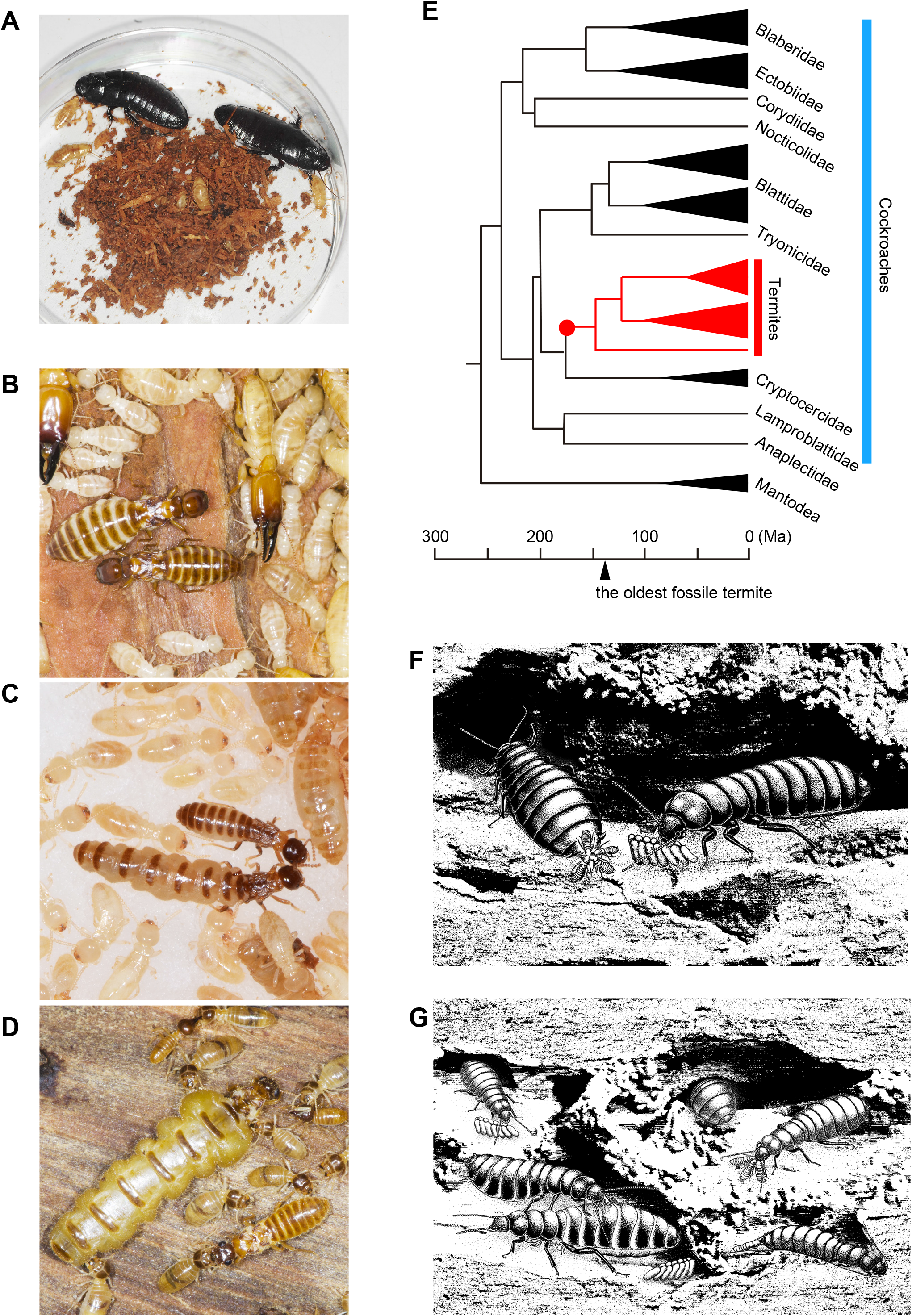
Termite eusociality originated from a lineage of subsocial woodroach. (*A*) Monogamy in a woodroach *Cryptocercus punctulatus*, which is an oviparous cockroach with biparental care. (*B*-*D*) A primary king (PK) and a primary queen (PQ) of a damp-wood termite *Zootermopsis nevadensis*, a subterranean termite *Reticulitermes speratus* and a higher termite *Nasutitermes takasagoensis*, respectively. (*E*) Phylogenetic placement of termites. Termites are a sister group of Cryptocercidae. (*F*, Imaginary illustration of the incipient colony (*F*) and a developed family (*G*) of the first termite. The founder female produces a packaged egg mass as in the basal termite *Mastotermes darwiniensis*. The first brood helpers contribute to the production of younger sibs through wood excavation, trophallaxis, allogrooming, nest hygiene and so on. Photos by K.M. Illustrations by H.O and K.M.

The evolutionary transition from subsociality to eusociality, where the older brood spend a prolonged period in the immature stages within the natal nest and eventually become lifelong workers that contribute to the production of younger brood, is an enigma (Korb, 2008; Thorne, 1997). Although the shift of brood care from parents to older offspring, i.e., helpers, allows the parents to engage more in reproduction, selection is still acting at the individual level rather than the colony level at the subsocial stage. Why did helpers delay and eventually sacrifice their own reproduction under individual-level selection? We show that this paradox can be solved by genomic imprinting.

Recent studies reveal that the transfer of epigenetic marks from parents to offspring can influence offspring phenotype independently of DNA sequence (Bonduriansky and Day, 2008; Chong and Whitelaw, 2004; Ferguson-Smith, 2011; Jablonka and Raz, 2009). Such ‘epigenetic inheritance’ provides the key to many unsolved puzzles in a wide range of biological processes by circumventing the limitations of genetic inheritance (Bonduriansky and Day, 2008; Matsuura, 2020a; Pál and Miklós, 1999). Genomic imprinting can be differentially marked in eggs and sperm, and inheritance of these epigenetic marks causes genes to be expressed in a parental-origin-specific manner in the offspring (Reik and Walter, 2001). Genomic imprinting has been proposed as a mechanism underlying the parental effects on caste determination (Matsuura et al., 2018) and the evolution of asexual queen succession in termites (Matsuura, 2017; Matsuura et al., 2018, 2009) (see supplementary text, Figs. S1 and S2). Thus, queen- and king-specific epigenetic marks antagonistically influence the development of offspring resulting in “neuter” workers whose sexual development is suppressed due to counterbalanced maternal and paternal imprinting.

## Model

### General framework

To consider the evolutionary transition from subsocial to eusocial in the ancestor of termites, we suppose a diploid subsocial organism in which a male and a female cooperatively found a new colony in an enclosed habitat. We performed evolutionary simulations by running colony generation *G*, where each generation includes a colony life cycle as a part of the simulation (Fig. 2, Fig. S6). The colony life cycle is given as follows. An individual has longevity *Φ* (years), where its age *i* varies from 0 to *Φ*. Individuals spend *τ* (≥ *τ*_*min*_: the shortest time to become alates) in their natal nests and become alates that disperse. The colony dies when the parents die. Thus, colony longevity *T* is given by *Φ* – *τ* (Fig. 2*A*), where colony age *t* varies from 0 to *T*. Fitness gain *ω*_*t*_ of the colony at *t* is the sum of the reproductive values of dispersing alates (Fig. 2*A*). Total fitness of the colony *W* is given by

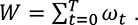

**Figure 2.**
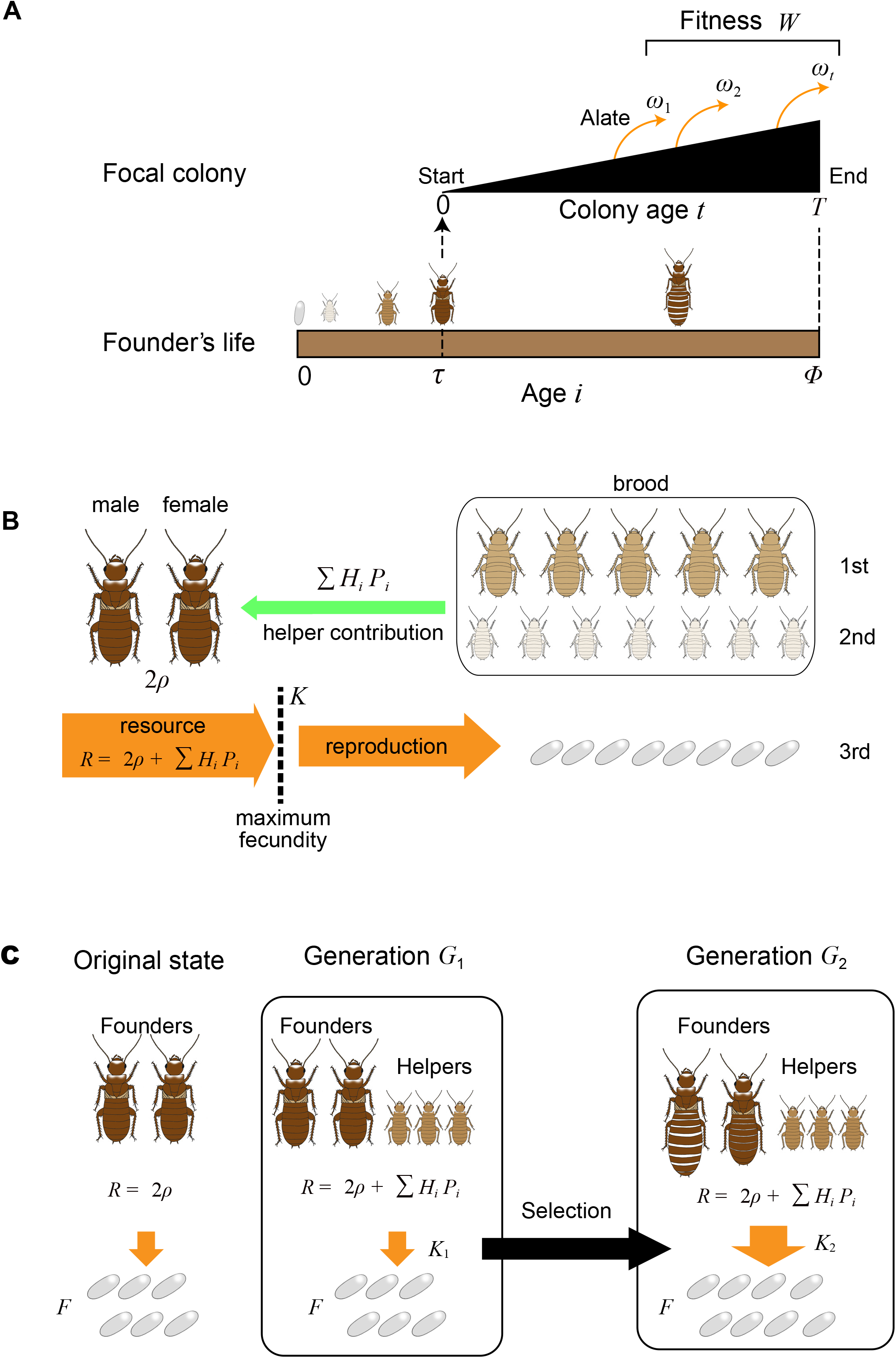
Schematic view of the model life cycle of a subsocial ancestor of termites just before the origin of eusociality. (*A*) Colony life cycle. Individuals become alates (imagoes) and leave their natal nest at age *τ*. They disperse and establish new colonies by pairing with an unrelated individual. The colony produces alates *ω*_*t*_ at colony age *t*. Total number of alates produced *W* during the colony lifespan (*Φ* – *τ*) is defined as the fitness of the focal colony. (*B*) Presence of older brood in the nest contributes to resource acquisition of parents. (*C*), Additional resources can be a selective force for the evolution of higher fecundity of founders. Without helpers, the founders reproduce by using their own resources 2*ρ* (left). Even if there were additional resources from the helpers’ contributions, the founders would not be able to use them for reproduction due to the upper limit of their fecundity *K*_1_ (center). Selection favors higher fecundity *K*_2_ of the founders so as to fully utilize the additional resources for reproduction (right). See also Figs. S5 and S6 for the detailed scheme. Illustrations by K.M.

### Colony foundation

We consider a monogamous pair of a male and a female founding a new colony in a nest inside wood. The work performance of each parent is *ρ* (J/year), that is, the amount of resources acquired by a parent. Both the male and female founder contribute to the production of eggs equally. The available resources for egg production of a *t*-year-old colony is *R*_*t*_. In the first year (0-year-old colony), the parents produce eggs by using the available resources *R*_0_ (J) (= 2*ρ*) (Fig. 2*B*). The number of first brood eggs *F*_0_ is given by

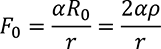

where *α* is the metabolic efficiency of the parents to convert the resources into eggs, *r* is the cost for producing one egg. For simplicity, we set *α*/*r* = 1, i.e., *F*_0_ = 2*ρ*. The final family composition of the 0-year-old colony is: a founding male, a founding female and first brood (*F*_0_ eggs).

Second year (1-year-old colony): The 1st brood develop into 1-year old larvae, which are too small and undeveloped to contribute to social labor. In the second year, the available resources *R*_1_ (J) are still 2*ρ*. Parents produce *F*_0_ eggs of the 2nd brood in the second year. Therefore, the final family composition of the 1-year-old colony is: a founding male, a founding female, first brood (*F*_0_ 1-year-old larva) and 2nd brood (*F*_0_ eggs).

In the third year, the 2nd brood are 1-year-old, which do not contribute to social labor. The 1st brood are 2-years-old, at which time they begin to contribute to social labor as helpers.

### Work performance of helpers

Work performance *P*_*i*_ (J/year) of a helper offspring increases with age *i*. Eggs (0-year-old) and 1-year-old larva do not contribute to social labor. Offspring older than 5 years achieve the maximum performance *β* · *ρ* in their lifetime, where *β* is the relative work efficiency of helpers compared to the work performance of a parent *ρ* (Figs. S3 and S4). If *β* is 1, mature helpers (offspring older than 5 years) have the same work performance as a parent. The 2-4 year old offspring have intermediate performance. Hence, the work performance *P*_*i*_ of a helper at age *i* (year) is given by

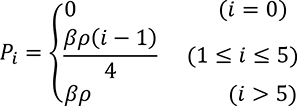

Total acquired resources *R*_*t*_ (J) are

*R*_*t*_ = 2*ρ* + ∑ *H*_*i*_ · *P*_*i*_,

where *H*_*i*_ is the number of *i*-year-old helpers.

### Reproductive capacity and genomic imprinting

Relationship between the amount of resources *R*_*t*_ and the number of eggs *F*_*t*_ produced in year *t* is given by

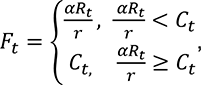

where *C*_*t*_ is the reproductive capacity of the parents at colony age *t* (Fig. 3*A*). Reproductive capacity of the parents *C*_*t*_ increases with their sexual development. Reproductive capacity of the parents *C*_*t*_ is given by

**Figure 3.**
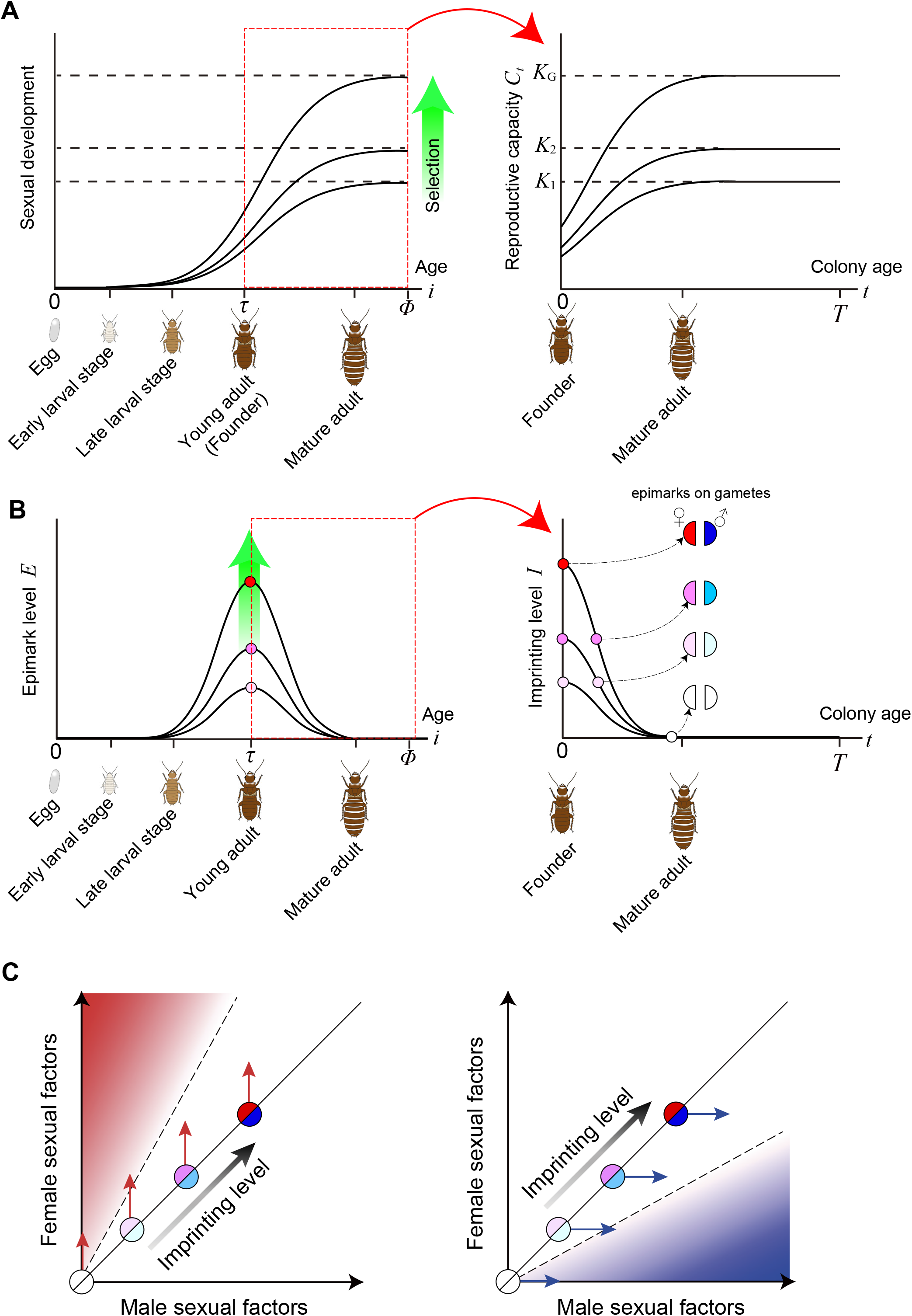
The genomic imprinting model for the origin of eusociality. Sex-antagonistic action of maternally- and paternally-inherited epimarks retains the offspring in the neuter state. (*A*) Sexual development of individuals (left) and reproductive capacity of founders *C*_*t*_ (right). (*B*) Development of epimarks in relation to sexual development of the parents (left) and unerased sex-specific epimarks that carryover to the next generation through gametes (right). Reproductive capacity of the parents *C*_*t*_ is acquired through canalization of the expression of sexual development genes through epigenetic modification *E*. Therefore, *E_t_* is expressed as the derivative of 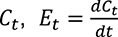. Blue and red semicircles indicate paternally- and maternally-inherited epimarks, respectively. Darkness of the color represents the strength of epimarks. (*C*) Antagonistic action of maternal and paternal imprinting in the offspring. Illustrations by K.M.

*C*_*t*_ = *K*_*G*_ · tanh (*t* + 1),

where *K*_*G*_ is the maximum fecundity of the parents (Fig. 3*A*).

The maximum reproductive capacity *K*_*G*_ is determined by sexual development of the parents, which is acquired through canalization of the expression of sexual development genes through epigenetic modification (epimark level) *E*_*t*_ (Fig. 3*B*).

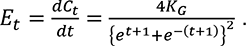

Epigenetic marks are partially inherited by offspring, which leads to a genomic imprinting effect on the sexual development of the offspring. Therefore, imprinting level *I*_*t*_ at colony age *t* is given by

*I*_*t*_ = *εE*_*t*_.

where *ε* is the transmission rate of parental epimarks to offspring (Fig. 3*B*). Antagonistic action of paternal imprinting *I*_*m*_ and maternal imprinting *I*_*f*_ delay the initiation of sexual development of offspring, resulting in prolongation of the offspring’s time to become alates *τ* (Fig. 3*C*). For simplicity, we assume

*I*_*m*_ = *I*_*f*_ = *I*_*t*_.

### Genomic imprinting and maturation delay of offspring

The time *τ* required to become alates, numerically calculated from *I*_*t*_ and its boundary conditions, is given by

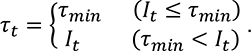

The offspring that have spent *τ*_*t*_ after hatching in year *t* become alates, contributing to the founders’ fitness (Fig. 2*A*). We set *τ*_*min*_ = 4 (years) based on the subsocial sister group *Cryptocercus* woodroaches, whose maturity takes about 6 years (Cleveland et al., 1934; Nalepa, 1984). Note that when an offspring stays longer in the colony as a helper, its direct contribution to the yearly fitness is not only delayed, but also its residual life span as a colony founder is shortened. However, its contribution as a helper will increase the production of future offspring (younger brothers and sisters) in the current colony (Fig. 2*B*). This trade-off between the contribution by staying as helpers and leaving success as alates is key for the evolution of eusociality.

### Colony fitness and selection process

Fitness gain *ω*_*G*,*t*_ of the colony at *t* is the sum of the reproductive values of dispersing alates with different ages *i*, such that

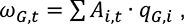

where *A*_*i*,*t*_ is the number of alates at age *i* produced in year *t* and *q*_*G*,*i*_ is the reproductive value of an alate at age *i*. In exchange for a prolonged helper period, the reproductive value of an alate *q*_*G*,*i*_ declines with the shortening of its residual life span (*Φ* − *i*) after colony foundation. This discounted reproductive value of an alate at age *i* is numerically calculated from the fitness function *ω*_*G*−1,*t*_ of the previous generation *G* − 1, such that

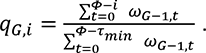

In order to calculate the fitness of the first generation *ω*_1,*t*_, we set *q*_1,*i*_ = 1 because there is no imprinting effects on offspring (all offspring reaching age *τ*_*min*_ become alates without delay). Total fitness of generation *G* is given by

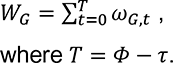

When the resources exceed the maximum reproductive capacity of the parents *K*_*G*_, there is no benefit to the colony since they cannot be used to produce additional eggs. The maximum amount of excess resources Δ*R* due to the limitation of reproductive capacity *K*_*G*_ is given by

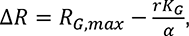

where *R*_*G*,*max*_ is the maximum yearly resources of a colony at generation *G*. Mutation from *K* to *K* + 1 takes place if 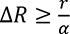 (at least one egg can be produced by the surplus resource) (Fig. 2*C*, Figs. S5 and S6). Then selection should fix this mutation if *W*_*G*_ ≥ *W*_*G*−1_ (fitness can be increased by mutation). Higher reproductive capacity results from increased epigenetic modification (i.e., stronger canalization (Rice et al., 2016)) of sexual development genes, which results in a higher imprinting level in offspring. Greater imprinting further prolongs the helper period resulting in in increased resources in the next generation *G* + 1, which again produces surplus resources Δ*R*. This whole process may be repeated to increase reproductive capacity *K*_*G*_ as long as the two above-mentioned conditions are satisfied (Figs. S5 and S6). We consider that eusociality arises at the time when the first-brood offspring lose all direct fitness. We also note the time at which the first-brood helpers spend their whole life in the colony, i.e., the appearance of lifelong helpers.

## Results

We show that genomic imprinting can explain the origin of eusociality, i.e., the first occurrence of a lifelong neuter caste. To consider the evolutionary transition from subsocial to eusocial in the ancestor of termites, we suppose a diploid subsocial organism in which a male and a female cooperatively found a new colony in an enclosed habitat (Figs. 1*F*, 1*G* and 2). We developed a genomic imprinting model that predicts the evolution of eusociality via natural selection when parental reproductive capacity is linked to a delay in the sexual development of offspring through transgenerational epigenetic marks that act in a sex-antagonistic manner. We performed evolutionary simulations by running colony generation, where each generation includes a colony life cycle as a part of the simulation (Fig. 2, Fig. S6). Presence of helpers contribute to social labor, which increases the resources available for egg production (Fig. 2*B*). Selection favors higher reproductive capacity of the parents so as to fully utilize the available resources (Fig. 2*C*). Acquisition of greater reproductive potential is derived from higher epigenetic modification (canalization) of the genes for sexual development (Fig. 3 *A* and *B*). This leads to higher genomic imprinting and thus to the prolonged helper period of the offspring (Fig. 3*C*).

Our numerical algorithm demonstrates that genomic imprinting can drive eusociality via natural selection on colony founders (Fig. 4). The evolutionary transition from subsocial to eusocial systems readily occurs when the epimark transmission rate *ε* and/or the helpers’ work performance *β* are high (Fig. 4*A*). Because helpers increase the resources available for egg production (Figs. S3 and S4), natural selection acts on the parents to increase production of alates. Greater reproductive potential requires higher levels of epigenetic modification. Trans-generational carry-over of sex-specific epimarks leads to higher genomic imprinting and thus to a prolonged helper period for offspring. This evolutionary feedback between natural selection and genomic imprinting facilitates the transition from subsociality to eusociality (Fig. 4*B*). At some generation, the first-brood offspring lose their direct fitness (the origin of eusociality, red arrowheads in Fig. 4 *B* and *C*), and then spend their entire life in the colony as helpers (open arrowheads in Fig. 4 *B* and *C*). Here the offspring are essentially the extended phenotype of the parents via genomic imprinting.

**Figure 4.**
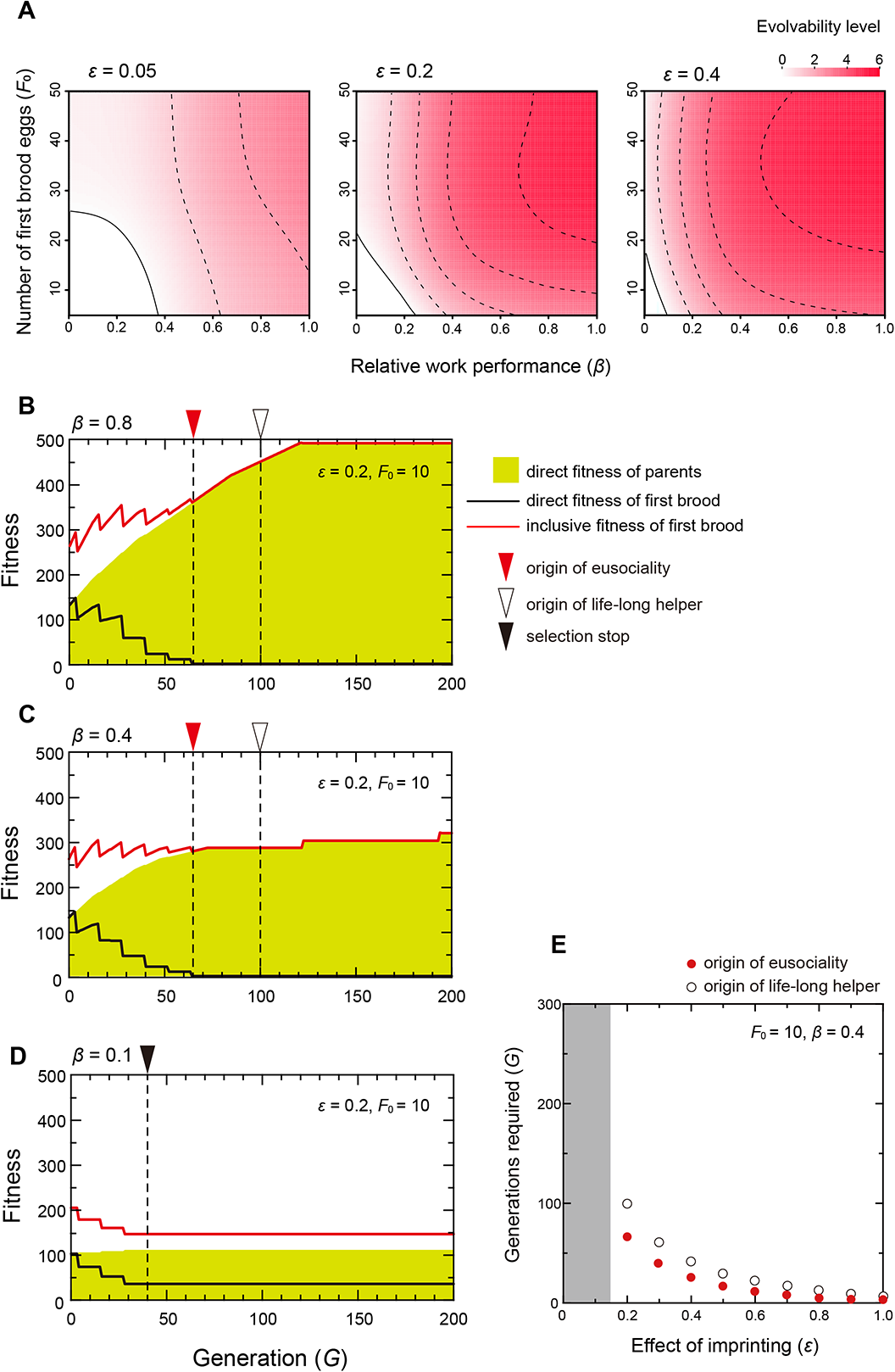
Evolvability of eusociality. (*A*) The effect of work performance of helpers *β* and initial fecundity of the parents *F*_0_ on the evolvability of eusociality. Eusociality is easily achieved under high epimark transmission rate *ε* and/or high work performance of helpers *β*. Evolvability level is evaluated by the level of eusociality acquisition (indicated by the intensity of red color) when colony longevity *T* is varied from 10 to 15 years (Level 0: no acquisition; level 6: acquisition under all *T* cases). Eusociality acquisition is defined as the appearance of a helper with no direct fitness (inclusive fitness only). (*B*-*D*) Fitness dynamics of the parents and the first brood offspring under different work performance levels of helpers *β* (*T* = 13, *ε* = 0.2, *F*_0_ = 10). Green area indicates the direct fitness of parents (the number of alates produced by a colony). Black and red lines indicate the direct and inclusive fitness of the first brood, respectively. Red arrowheads indicate the origin of eusociality (the appearance of a helper with no direct fitness). Open arrowheads indicate the appearance of a lifelong helper. Black arrowheads indicate the point when selection stops, i.e., the direct fitness of the parents decreases from that of the previous generation in numerical calculation. (*E*) Effect of genomic imprinting level *I*_*t*_ (i.e., *ε* for a given *E*_*t*_) on the generation required to reach eusociality. Red and open circles indicate the origin of eusociality and the appearance of a lifelong helper, respectively. Gray area indicates the region where eusociality is not achieved.

This feedback process generating eusociality creates parent-offspring conflict frequently because the first brood decreases its direct (and even inclusive) fitness by prolonged helper periods (zigzag black and red lines in Fig 4*B*), but eventually increases its inclusive fitness. Eusociality can still be achieved even if the helpers’ work performance is reduced by one-half (*β* = 0.8 → 0.4), although increase in the rate of parental direct fitness is slowed down (Fig. 4*C*). When the helpers’ work performance is further reduced (*β* = 0.1), natural selection does not drive eusociality (Fig. 4*D*), because helpers cannot produce sufficient resources for increased egg production. Most importantly, the level of genomic imprinting is a key factor driving eusociality; the number of generations required for eusociality (or lifelong helper) to evolve sharply decreases with imprinting strength (i.e., epimark transmission rate *ε* for a given *E*_*t*_) (Fig. 4*E*). Eusociality cannot be achieved if imprinting strength is too low (gray area in Fig. 4*E*). Also, the development of eusociality is impossible if colony longevity *T* is too short (Fig. S7), because the contribution of the first brood helpers cannot be used for the production of alates in the colony. It should be noted that when *T* is larger, it takes more time for eusociality to evolve. This is because it takes more generations for the first brood to become lifelong helpers when they have a longer lifespan (*Φ* = *T* + *τ*_*min*_).

## Discussion

The theoretical framework of the present model assumes iteroparity of the founder parents within the nest. This assumption is reasonable because all eusocial insects are iteroparous, which is essential to produce staggered age classes within colonies, enabling older offspring to rear younger siblings. In contrast, the *Cryptocercus* woodroach (the sister group of termites) is semelparous (Nalepa, 1988) and not eusocial. Without iteroparity (no second nor later brood), the evolutionary feedback loop cannot be established. Unlike the Hymenoptera where eusociality evolved multiple times, eusociality evolved only once in the cockroaches (termites) around 150 Myr ago (Bourguignon et al., 2015; Legendre et al., 2015). This single origin of eusociality in Blattodea is likely due to the fact that the acquisition of within-nest iteroparity occurred only once in this taxon, although non-nesting and serial nesting iteroparity are known in some species (Trumbo, 2013).

The developmental time to reach maturity should have been long in the subsocial termite ancestor due to a nitrogen-poor wood diet as in wood-feeding cockroaches in which developmental time ranges from three to seven years (Nalepa, 2015). Among wood-feeding cockroaches, the sister-group of termites, *Cryptocercus*, is the only known oviparous cockroach with biparental care and altricial development (Bell et al., 2007; Klass et al., 2008) under biparental care, crucial conditions for the origin of eusociality (Nalepa, 2010). The prolonged altricial period of the offspring creates overlapping broods within a colony providing the opportunity for older brood to help younger sibs (Thorne, 1997). We can reasonably expect that in the termite ancestor the presence of older brood was advantageous for the parents and younger sibs because they provided nest excavation, trophallaxis, allogrooming, defense and other forms of brood care. Therefore, we can expect high work performance *β* of the helpers in the subsocial ancestral state. In our model, a certain level of work performance *β* is necessary for the origin of eusociality. Interestingly, however, *β* can be as low as 0.1 when other conditions are favorable (e.g., *T* ≥ 15, *ε* ≥ 0.2, *F*_0_ ≥ 20, Fig. 4*A*).

As we discussed at the beginning, the evolutionary mechanisms driving the origin of eusociality and selection for the maintenance and later elaboration of eusociality should be discussed separately. We demonstrated that the acquisition of eusociality promotes the production of alates by a colony and also increases the inclusive fitness of the first brood compared with the initial subsocial state (Fig. 4 *B* and *C*). In this respect, our findings are consistent with inclusive fitness theory. Most importantly, however, we found that inclusive fitness theory itself cannot explain the origin of eusociality. During the transition from the subsocial to the eusocial state, inclusive fitness of the first brood results in a zigzag line alternating up and down for levels of fitness (Fig. 4 *B* and *C*), because there is a time lag between the origin of a prolonged helper period in the first brood offspring and an increase in the reproductive capacity of the parents to utilize the additional resources. This indicates that selection for higher inclusive fitness of the offspring cannot get over the selection valley (it goes down and then up again) and thus cannot generate lifelong helpers.

The earlier hypothesis that tried to overcome this selection valley is parental manipulation (Alexander, 1974): parents restrict the reproductive options of some offspring so that they assist in the production of fully fertile offspring. A serious problem with this hypothesis is that counter-selection on the offspring would make them overcome parental manipulation. Moreover, this hypothesis also cannot explain the origin of eusociality, although parental manipulation may increase the parental fitness and the inclusive fitness of the sterile workers in a eusocial state. Selection cannot act on the parents *a priori* in favor of restricting offspring’s reproductive development in a subsocial state because it merely reduces the number of fertile offspring and thus their own fitness unless it can be coupled with the subsequent evolution of the ability of the parents to utilize the additional resources for reproduction. A related idea of parental manipulation is the subfertility hypothesis of West-Eberhard (West-Eberhard, 1975), which assumes that subfertile offspring can be produced by inescapable factors such as food shortage and these subfertile offspring would give up reproduction and accept a worker role.

Genomic imprinting can be considered one such inescapable factor that produces subfertile offspring. In our genomic imprinting model, natural selection acts on the reproductive capacity of parents so as to increase their direct fitness. Greater reproductive capacity requires higher epigenetic modification of the genes for sexual development (Rice et al., 2016). Sex-specific epigenetic modifications are potential mechanisms underlying sexual dimorphism in various taxa (Hall et al., 2014; Pennell and Morrow, 2013). When sex-specific epimarks that canalize sexually dimorphic development are inherited by the offspring (Rice et al., 2012), sex-antagonistic action (Gadagkar, 2000) of the maternally and paternally inherited epimarks generates the delay of sex expression in the offspring, i.e., the first brood helpers. The offspring’s phenotype resulting from the transgenerational epigenetic inheritance is an extended phenotype of the parents, and is thus inescapable for the offspring.

Ultimately, whether eusociality spreads in a population depends on the specific benefits of altruistic behavior of the sterile workers relative to their own independent reproduction (Hamilton, 1964). Right at and after the origin of eusociality, the fitness interests of the parents and the offspring coincide with each other (Fig. 4 *B* and *C*, Fig. S7): the inclusive (i.e., indirect) fitness of the sterile helpers coincides with the direct fitness of the parents (i.e., colony fitness). From this point on, the subsequent evolution of eusociality is promoted by kin selection, where Hamilton’s rule *r* > *c*/*b* is satisfied. In the current numerical analysis, relatedness *r* is 0.5 because we assume lifelong monogamy of the parents (Boomsma, 2009), and the work performance of helpers *β* corresponds to the indirect benefit *b*, while the fitness loss produced by the prolonged helper period *τ* is the cost *c* of helpers. Then kin selection (i.e., colony level selection) drives the evolution of extreme morphology and self-sacrificing behavior of soldiers highly specialized in their defensive tasks (Deligne et al., 1981; Šobotník et al., 2012), sophisticated pheromonal communication regulating caste differentiation (Matsumura et al., 1968; Matsuura et al., 2010, 2007; Mitaka et al., 2017), and loss of totipotency, all of which accelerate the reproductive division of labor.

This genomic imprinting theory is based on the fact that genomic imprinting operates in caste determination of termites, where sexual development of the offspring is influenced by the sexual phenotypes of parents via transgenerational epigenetic effects (Matsuura, 2020b; Matsuura et al., 2018). Many molecular agents are known to mediate genomic imprinting such as DNA methylation, histone modifications, noncoding RNAs and transcription factors (Ferguson-Smith, 2011; Reik and Walter, 2001). Sex-specific methylation is a potential mechanism underlying sexual dimorphism in various taxa (Hall et al., 2014; Pennell and Morrow, 2013), and a recent molecular study demonstrated that DNA methylation regulates female fecundity in insects (Zhang et al., 2015). In termites, the level of DNA methylation is known to be considerably higher than in any other social insect studied to date (Glastad et al., 2016). Future studies are needed to determine the exact molecular mechanism underlying the genomic imprinting of termites, even though our conclusion holds irrespective of the exact agents.

The current theory to explain the origin of eusociality has the potential for broad applications to the origin of sterile workers in various diploid animals such as the ambrosia beetle *Austroplatypus incompertus* (Smith et al., 2018) and the naked mole-rat *Heterocephalus glaber* (Jarvis, 1981). It remains to be investigated in the future whether this theory can be extended to the origin of eusocial Hymenoptera, in which the existence and the role of genomic imprinting in the current caste system are still controversial. This study shows that transgenerational epigenetic inheritance can play a key role in the inception of eusociality and provides a novel framework for studying the evolution of social systems in various organisms.

### Data Availability

Major simulation data supporting the findings of this study are available within the paper and Supplementary Information. Simulation data can be generated within the custom-made code and the parameter sets provided. Custom-made simulation code is available via GitHub at (https://github.com/ItoHiromu/Eusoc-Sim).

## Acknowledgments.

We thank N.E. Pierce, Y. Iwasa, H. Saze, S. Gavrilets, K. Tsuji, E. Tasaki, T. Ishibashi, N. Mizumoto and M. Takata for discussions of this topic. K.M. acknowledge financial support from JSPS through grant 18H05268, 18H05372 and 25221206. E.L.V. was supported by the Urban Entomology Endowment at Texas A&M University.

## Supplementary Information

### Backgrounds

We here introduce the latest studies on the transgenerational epigenetic inheritance and genomic imprinting in termites following the recent review by Matsuura (Matsuura, 2020).

#### 1. Transgenerational epigenetic inheritance; a paradigm shift in biology

Evolution by natural selection requires phenotypic variation, inheritance and selection forces. Charles Darwin conceived the theory of natural selection without knowing the agent of inheritance. Biology is traditionally dominated by the idea of genetic determinism, which is focused almost exclusively on genetic inheritance and processes that change gene frequencies. Recent empirical studies have convincingly demonstrated transgenerational epigenetic inheritance in various eukaryotic organisms including yeasts (Grewal and Klar, 1996), plants (Hauser et al., 2011; Miryeganeh and Saze, 2020) nematodes (Remy, 2010; Serobyan and Sommer, 2017), mammals (Daxinger and Whitelaw, 2012; Perez and Lehner, 2019) and insects (Mukherjee and Vilcinskas, 2019). The transfer of epigenetic marks from parents to offspring can influence offspring phenotype independent of the DNA sequence (Bonduriansky and Day, 2008). The concept of epigenetic inheritance provides solutions for many previously-unsolved puzzles by circumventing the limitation of genetic inheritance (Bonduriansky and Day, 2008; Jablonka and Raz, 2009; Szyf, 2014). Theoretical studies indicate that epigenetic inheritance is an important factor in evolution that can produce outcomes that are not anticipated under traditional genetics. Therefore, it is apparent that biology is in a major transitional period that will take it forward from the previously narrow perspective.

Over the past decade there has been a rapid accumulation of definitive evidence for the molecular mechanisms underlying transgenerational epigenetic inheritance (Greer and Shi, 2012; Liberman et al., 2019; Martin and Zhang, 2007; Miska and Ferguson-smith, 2016; Moazed, 2011). In principle, any molecular change in the zygote other than alterations in DNA sequence could carry non-genetic information including DNA methylation, histone modification, non-coding RNA, prions and microbiota.

#### 2. Genomic imprinting as a carryover effect of parental epimarks

The effects of transgenerational epigenetic inheritance on offspring are often sex-specific (Bonduriansky and Day, 2008; Vigé et al., 2008). A phenomenon related to transgenerational epigenetic inheritance is genomic imprinting, whereby epimark transmission depends on parental sex, resulting in parent-of-origin allelic expressions (Gregg et al., 2010; Lawson et al., 2013; Xu et al., 2014). Recent evidence also suggests that imprinting effects can depend on offspring sex (Hager et al., 2008).

Kinship theory of genomic imprinting focuses on a conflict (Haig, 2014) between the maternal and paternal genes (see also Holman & Kokko (Holman and Kokko, 2014) and Patten et al. (Patten et al., 2014)). Genomic imprinting can occur as a by-product of another advantageous process such as retrotransposon silencing (Suzuki et al., 2007) or canalization of sexually dimorphic development (Rice et al., 2012) even if imprinting itself has no fitness advantage. The most parsimonious mechanistic explanation for the origin of genomic imprinting would be incomplete erasure of epigenetic modifications, i.e., carryover effects of parental epimarks (Chong and Whitelaw, 2004; Kearns et al., 2000; Rice et al., 2016). Although epigenetic modifications acquired by parents are mostly erased during gametogenesis, which allows reestablishment of epigenetic marks in the next generation, a portion of epigenetic modifications can be inherited through the germline in a wide variety of taxa including mammals (Anway et al., 2005; Morgan and Bale, 2011), insects (Cavalli and Paro, 1998), yeast (Grewal and Klar, 1996) and plants (Miryeganeh and Saze, 2020).

The degree of sexual dimorphism will be associated with the degree of sex-specific epigenetic modifications (Deegan and Engel, 2019). Selection favoring a higher degree of sexual dimorphism would lead to a higher imprinting level, if the efficiency of erasure is not changed. This carryover effect of sex-specific parental epimarks on the offspring’s development can contribute to the determination of the phenotype (Matsuura et al., 2018; Rice et al., 2013, 2012).

#### 3. Asexual queen succession in termites

Transgenerational epigenetic inheritance and genomic imprinting have been attracting more and more attention in the study of insects because many phenomena involving non-genetic inheritance have been found in a number of insect species (Galbraith et al., 2016; Gegner et al., 2019; Matsuura, 2020; Mukherjee and Vilcinskas, 2019). In particular, the effects of parental phenotypes on the caste fate of offspring have been discovered in termites, which provides definitive evidences for non-genetic inheritance (Matsuura et al., 2018). A unique reproductive system named Asexual Queen Succession (AQS) has been identified in several termite species, in which workers, soldiers and alates are produced sexually while neotenic queens arise through thelytokous parthenogenesis and eventually replace the old queens (Hellemans and Roisin, 2020; Matsuura, 2017; Matsuura et al., 2009) (Fig. S1). Then, neotenic queens are replaced by subsequent cohorts of asexually-produced neotenic queens. Therefore, as long as the colonies survive, the queens are genetically immortal. This is advantageous for both the queens and other colony members because queens can boost reproduction without inbreeding.

After the first report of AQS in the Japanese subterranean termite *R. speratus* (Matsuura et al., 2009), AQS has been continuously found in other *Reticulitermes* species in other continents: *R. virginicus* in the US (Vargo et al., 2012) and *R. lucifugus* in Italy (Luchetti et al., 2013). Time-scaled phylogeny of the genus *Reticulitermes* demonstrated that AQS was absent in the basal lineage but has evolved at least three times independently, in East Asia (ca. 7-5 million years ago), in West Europe (ca. 10-5 million years ago) and in North America (less than 14.1 million years ago) (Dedeine et al., 2016). Recent surveys of the breeding systems of higher termites (Termitidae) in French Guiana have repeatedly identified AQS in Neotropical termitids including *Embiratermes neotenicus* (Syntermitinae) (Fougeyrollas et al., 2015), the humivorous termite *Cavitermes tuberosus* (Termitinae) (Fournier et al., 2016), *Silvestritermes minutus* (Syntermitinae) (Fougeyrollas et al., 2017) and *Palmitermes impostor* (Termitinae) (Hellemans et al., 2019). In all of these seven AQS species, the asexual neotenic queens derive from nymphs, i.e., the sexual pathway.

Why has AQS evolved repeatedly in phylogenetically distant lineages? The origin of AQS requires the simultaneous evolution of two distinct traits: parthenogenetic capacity and the developmental priority for parthenogenetic daughters to develop into secondary queens. Then a question emerges as to why parthenogenetic daughters and sexually-produced daughters differ in their developmental propensity. Nozaki et al. (2018) asked if in a non-AQS species, *R. okinawanus*, the female offspring produced by tychoparthenogenesis (i.e., occasional parthenogenesis) differentiate into secondary queens (Nozaki et al., 2018). Interestingly a significantly higher proportion of parthenogenetic daughters developed into neotenic queens (nymphoid queens) than sexually produced females (Nozaki et al., 2018). This suggest that the developmental propensity of parthenogenetic daughters to become neotenic queens existed prior to the origin of AQS, which could explain why AQS evolved repeatedly in phylogenetically distant lineages.

#### 4. Genomic imprinting in termites

Influence of parental phenotypes on the caste fate of offspring is known in the genus *Reticulitermes* (Hayashi et al., 2007; Kitade et al., 2010). For instance, the daughters produced by queens’ parthenogenesis and the daughters produced by the mating of nymph-derived queens and worker-derived males develop exclusively into nymphs. On the other hand, most of the daughters produced by the mating of worker-derived queens and nymph-derived males develop into workers. The theory of genomic imprinting explains the effects of parental phenotypes on caste fate of the offspring(Matsuura et al., 2018). Its prediction matches exactly with empirical data from both lab and field studies.

As shown in the scheme (Fig. S2), the theory of genomic imprinting consists of two stages: 1) development of phenotype-specific parental epimarks and 2) influence of parental imprinting on the development of offspring. Nymph-derived reproductives obtain higher levels of epimarks so as to canalize expression of sexual characters than ergatoids (worker-derived reproductives) (Fig. S2*A*). Then the maternally- and paternally-inherited epimarks influence the relative expression of growth regulatory genes (εg: growth factors) to sex regulatory genes (εs: sexual factors), and thus determine the caste fate of offspring (Fig. S2*B*). Growth with sexual development directs individuals to the sexual pathway (nymphs), while growth without sexual development restricts individuals to the neuter caste (workers). The double maternal imprinting of parthenogenetic daughters leads them to the sexual pathway in AQS species.

The predictions of genomic imprinting model match well both with the caste differentiation patterns of the AQS species *R. speratus* (Hayashi et al., 2007) (Fig. S2*B*) and those of the non-AQS species *R. kanmonensis*, *R. yaeyamanus* and *R. okinawanus* (Kitade et al., 2010). Genomic imprinting clearly explains why the daughters carrying only maternal chromosomes have the developmental propensity to become neotenic queens rather than daughters carrying both paternal and maternal chromosomes.

Genomic imprinting explains why parthenogens carrying only maternal chromosomes pre-adaptively have the epigenetic and thus developmental advantage to become neotenic queens (Matsuura et al., 2018; Nozaki et al., 2018). Many factors are known to mediate genomic imprinting such as DNA methylation, histone modifications, noncoding RNAs and transcription factors (Ferguson-Smith, 2011; Reik and Walter, 2001). The DNA methylation levels in termites are considerably higher than other hemimetabolous insects (Harrison et al., 2018). Moreover, termites shows caste- and sex-specific DNA methylation patterns (Glastad et al., 2016). The DNA methyltransferases DNMT1 and DNMT3 are known to be involved in maintenance of and mediating *de novo* DNA methylation, respectively (Goll and Bestor, 2005; Yan et al., 2015). RNA-seq analysis of *R. speratus* revealed sex- and caste-dependent expression patterns of the DNA methyltransferases, where alates have higher expression of *Dnmt1* than workers in females but not in males. Alates also showed significantly higher expression of *Dnmt3* than workers in males but not in female (Mitaka et al., 2020). These results seem to support the theory of genomic imprinting.

#### 5. Additional model explanation

##### Work performance of helpers

Within-nest iteroparity creates the overlapping broods within a colony, which provides the opportunity for older brood to contribute to the production and growth of younger sibs (Thorne, 1997). The presence of older brood is advantageous for the parents and younger sibs rather than imposing a deleterious effect because they provide nest excavation, trophallaxis, allogrooming, defense and other forms of brood care. Therefore, we can reasonably expect a high work performance *β* of the helpers in the subsocial ancestral state.

Work performance *P*_*i*_ (J/year) of a helper offspring increases with age *i*. Eggs (0-year-old) and 1-year-old larvae cannot contribute to social labor. Offspring older than 5-years achieve their maximum possible performance *β* · *ρ*, where *β* is the relative work efficiency of helpers compared to the work performance of a parent *ρ*. If *β* is 1, mature helpers have the same work performance as a parent. 2-4 year old offspring have intermediate performance. Thus the helper performance *P*_*i*_ of a helper at age *i* (year) is given by (Fig. S3)

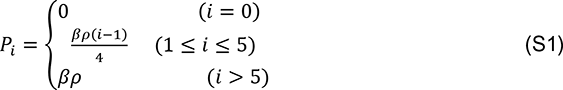

We also investigated the effect of the curvatures *s* of age-dependent increase in work performance *P*_*i*_ on the origin of eusociality (Fig. S4),

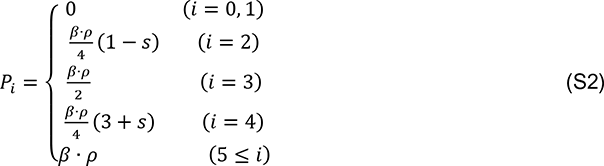

There is no qualitative change in the outcome when the curvatures *s* varies from 0 to 1 (the results were omitted).

---

**Fig. S1.**
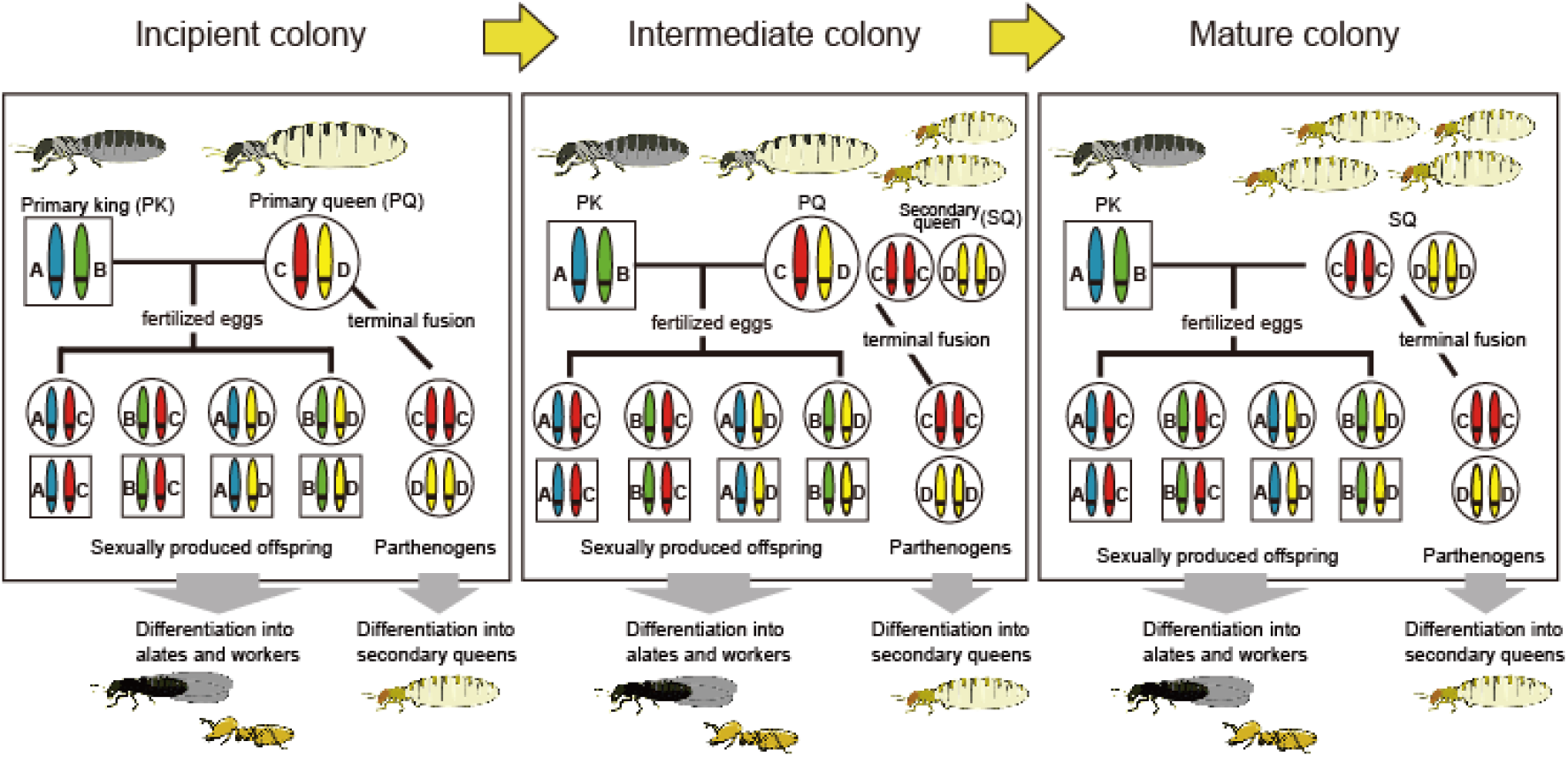
Asexual queen succession in termites. Secondary queens produced asexually by the primary queen differentiate within the colony and supplement egg production, eventually, replacing the primary queen (Matsuura, 2017; Matsuura et al., 2009). This breeding system enables the primary queen to maintain her full genetic contribution to the next generation, while avoiding any loss in genetic diversity from inbreeding.

**Fig. S2.**
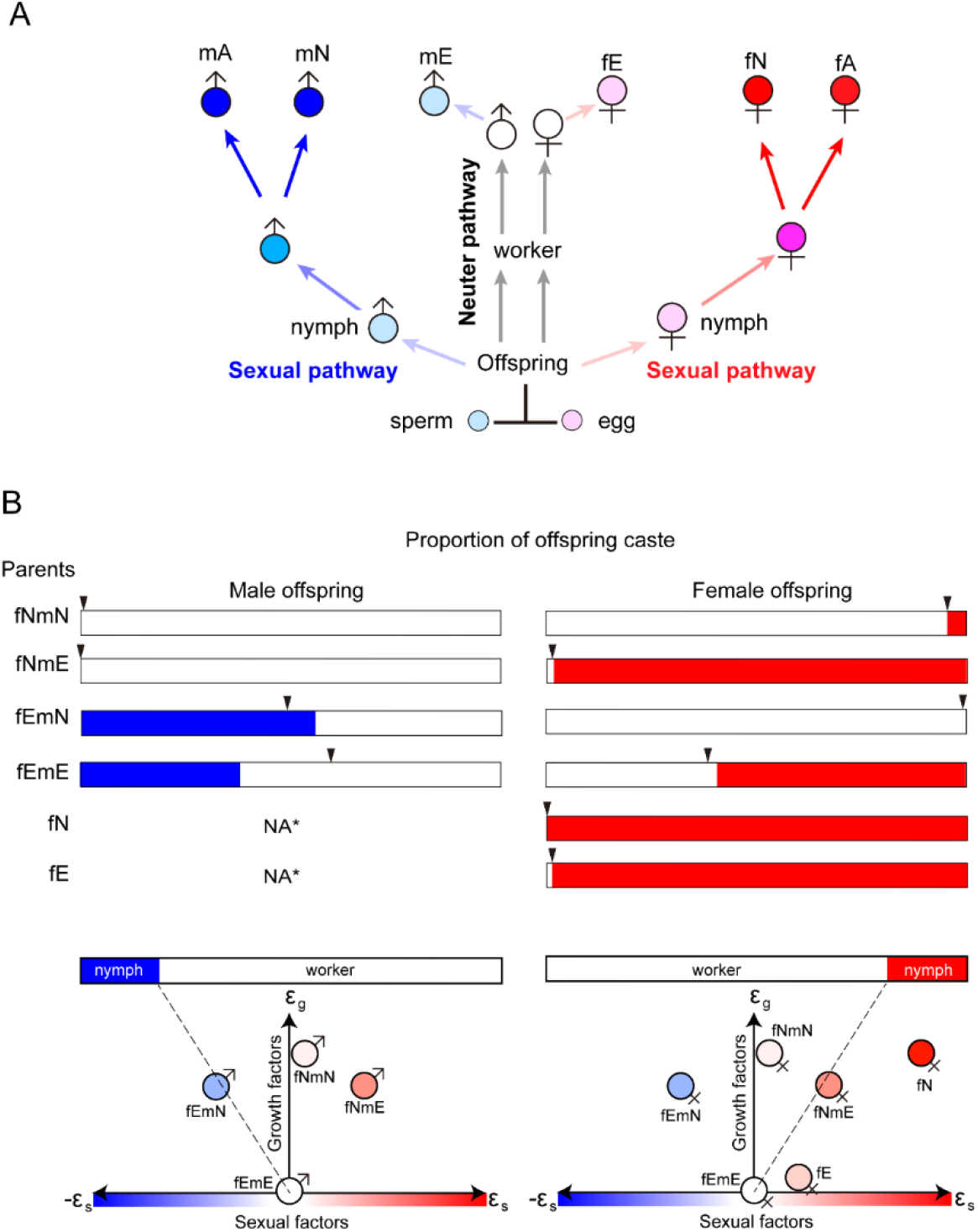
Genomic imprinting and caste determination in termites. (*A*) Caste differentiation and establishment of sex-specific epimarks. Alates (male: mA; female: fA) and nymphoids (male: mN; female: fN) develop with the full canalization of sexual development, while ergatoids (male: mE; female: fE) develop by a single sexual molt. Blue and red arrows indicate the development of male- and female-specific epimarks, respectively, and darkness of the color represents the strength of epimarks. (*B*) Combination of parental phenotypes and outcome of offspring caste (e.g., fEmN: female ergatoid and male nymphoid). The caste differentiation patterns predicted by the genomic imprinting model (bar graphs) and the empirical results (arrowheads) of *R. speratus* (Hayashi et al., 2007). *Parthenogenesis produces only female offspring. Expression levels of growth regulatory genes (εg) and sexual regulatory genes (εs) of the male- and female-offspring. Modified from Matsuura et al. (Matsuura et al., 2018).

**Fig. S3.**
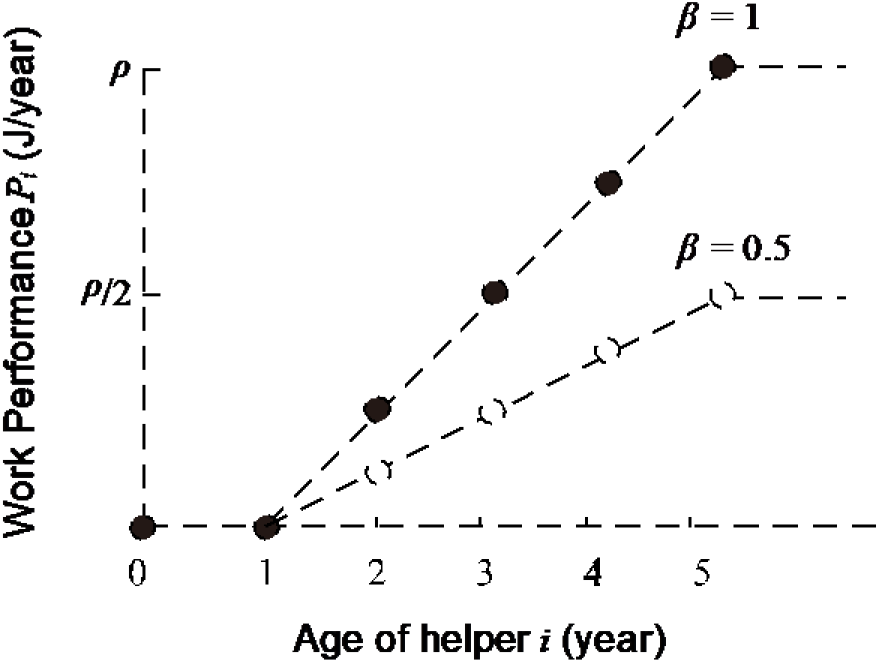
Relationship between age and work performance of helpers. Work performance *P*_*i*_ (J/year) of a helper increases with age *i* (year), where 0-year-old and 1-year-old helpers do not contribute to social labor. Offspring older than 5 years achieve their maximum possible performance *β* · *ρ*, where *β* is the relative work efficiency of helpers to the work performance of a parent *ρ*. If *β* is 1, mature helpers (offspring older than 5 years) have the same work performance as a parent (closed circles). The 2-4 year old offspring have intermediate performance levels. Hence, the work performance *P*_*i*_ of a helper at age *i* (year) is given by Equation (S1).

**Fig. S4.**
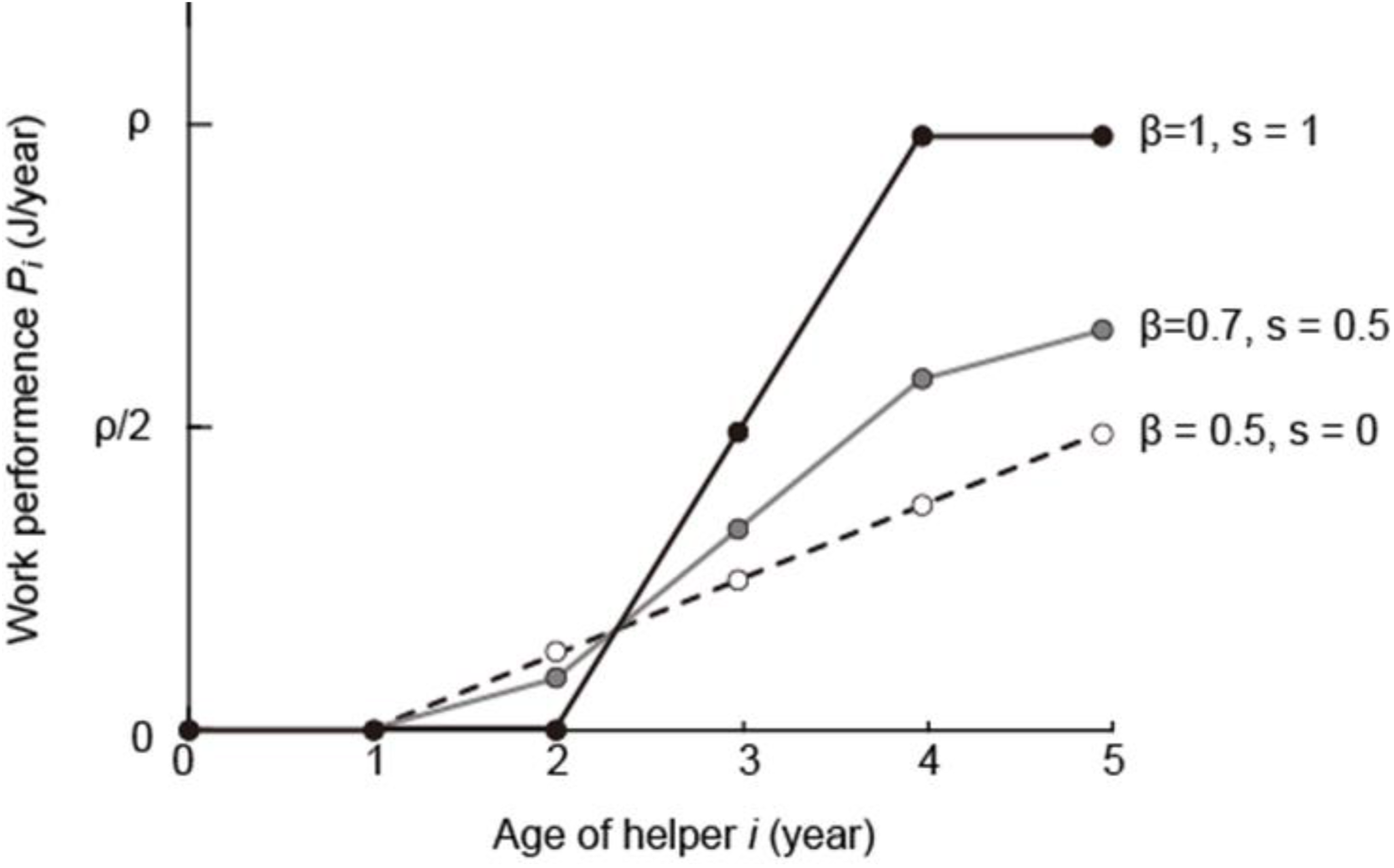
Relationship between age and work performance of helpers. The work performance *P*_*i*_ of a helper at age *i* (year) is given by Equation (S2).

**Fig. S5.**
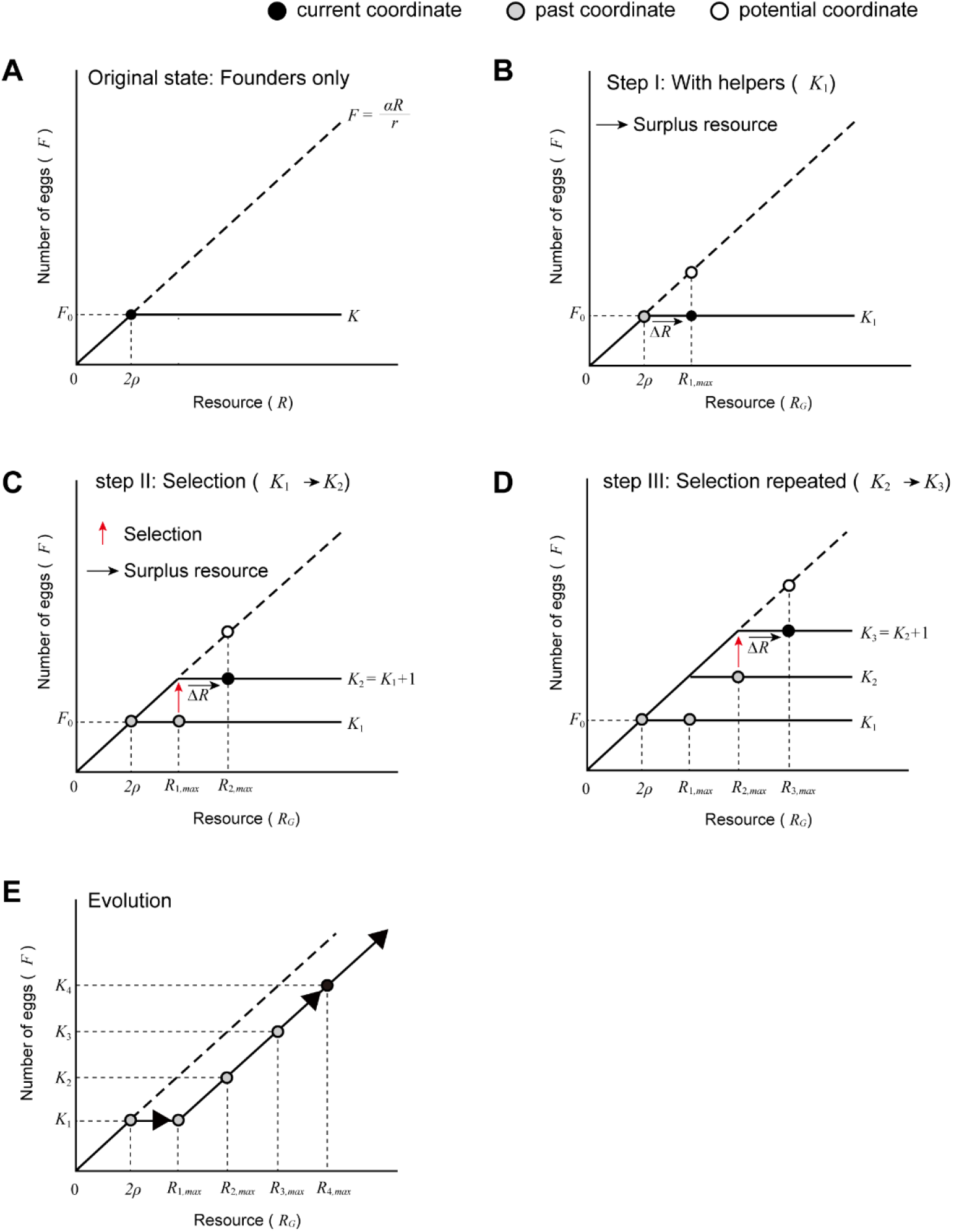
Selection scheme. (*A*) Original state. Without helpers, the founders reproduce by using their own resources 2*ρ*. A monogamous pair of a male and a female produce *F*_0_ eggs using their own resources 2*ρ*. (*B*) Step I (generation *G*_1_). Presence of helpers contribute to social labor, which increases the resources available Δ*R* for egg production. Nevertheless, the parents can produce only *F*_0_ eggs because of limitations on maximum reproductive capacity *K*_1_. (*C*) Step II (generation *G*_2_). Selection favors higher reproductive capacity of the parents so as to fully utilize the available resources (*K*_1_ → *K*_2_). Acquisition of greater reproductive potential is derived from higher epigenetic modification (canalization) of the genes for sexual development^30,31^. This leads to higher genomic imprinting and thus to the prolonged helper period of the offspring. Therefore, selection for higher *K* moves the coordinate from (*R*_1,*max*_, *K*_1_) to (*R*_2,*max*_, *K*_2_). (*D*) Step III (generation *G*_3_). In the same way as step II, selection favoring higher *K* to utilize the surplus resources moves the coordinate from (*R*_2,*max*_, *K*_2_) to (*R*_3,*max*_, *K*_3_). (*E*) The evolutionary process. Trace of the evolution of reproductive capacity starting from *K*_1_ (= *F*_0_).

**Fig. S6.**
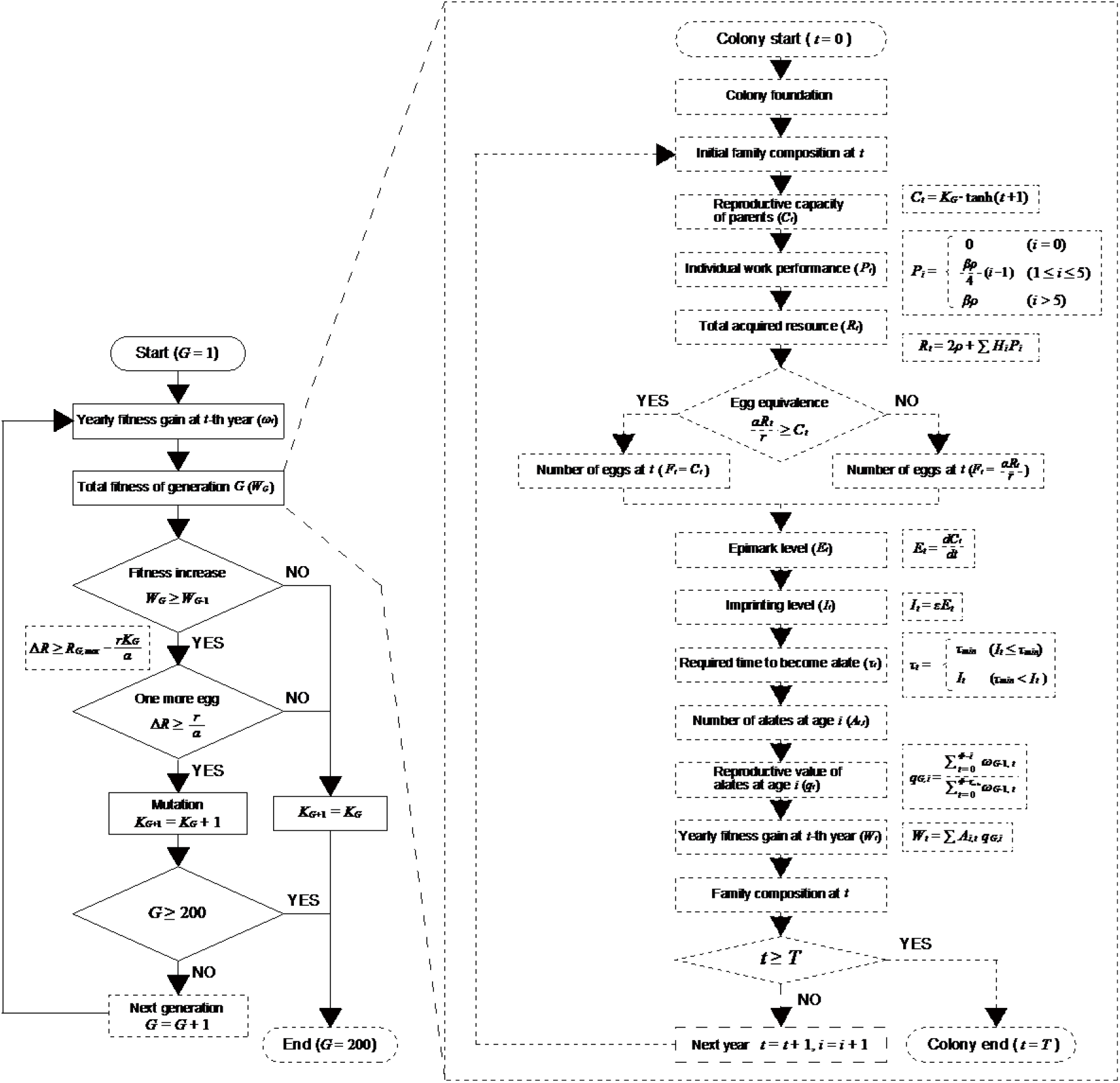
Simulation flow. See Model for the detailed procedure.

**Fig. S7.**
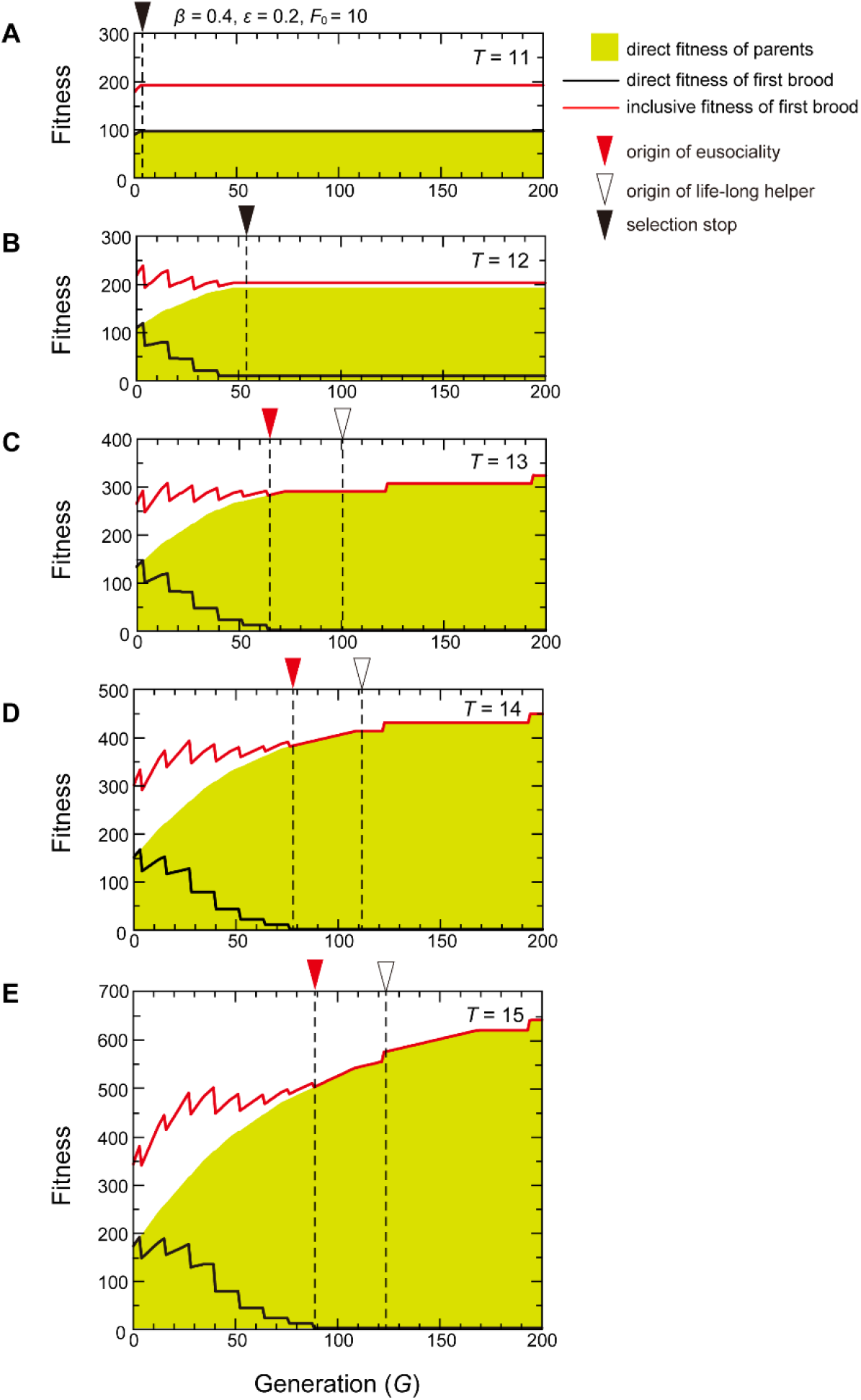
Colony longevity and evolvability of eusociality. Fitness dynamics of the parents and the first brood offspring under different values of colony longevity *T*. (*β* = 0.4, *ε* = 0.2, *F*_0_ = 10). (*A, B*) Colony lifespan is too short for the inception of eusociality. Selection stops, that is, the direct fitness of the parents decreases, at generation *G* = 5 when *T* = 11 (*A*) and at *G* = 53 when *T* = 12 (A) without reaching the threshold for eusociality. (*C-E*) Colony lifespan is sufficient for the evolution of eusociality. Eusociality originates at *G* = 65 when *T* = 13 (*C*), at *G* = 77 when *T* = 14 (D) and at *G* = 89 when *T* = 15 (*E*). Green area indicates the direct fitness of parents (the number of alates produced by a colony). Black and red lines indicate the direct and inclusive fitness of the first brood, respectively. Red arrowheads indicate the origin of eusociality (the appearance of a helper with no direct fitness). Open arrowheads indicate the appearance of lifelong helper. Black arrowheads indicate the point when selection stops, i.e., the direct fitness of the parents is lower than that of the previous generation in numerical calculation.

**Table S1.** The parameters of the genomic imprinting model.

| Symbol | Definition |
| --- | --- |
| $i$ | Age of offspring (years) |
| $\Phi$ | Longevity of individuals (years) |
| $\tau$ | Time from birth to dispersal (years) |
| $t$ | Colony age (year), i.e., time after colony foundation |
| $T$ | Longevity of a colony, i.e., $\Phi - \tau$ (years) |
| $\omega_t$ | Fitness gain of the colony at $t$ |
| $W$ | Total fitness of a colony |
| $\rho$ | Work performance of an adult (a founding male or a founding female) |
| $R_t$ | The available resources for egg production of a $t$-year-old colony |
| $\alpha$ | Metabolic efficiency of the parents to convert the resources into eggs |
| $r$ | Cost of producing one egg |
| $F_t$ | The number of eggs produced in year $t$ |
| $P_i$ | Work performance (J/year), i.e., the total resources (J) acquired by a single helper at age $i$ in one year |
| $\beta$ | Relative work performance of a helper to that of a parent $\rho$ |
| $H_i$ | Number of $i$-year-old helpers |
| $C_t$ | Reproductive capacity (eggs/year) of the parents of a $t$-year-old colony, that is, the maximum number of eggs produced in year $t$ |
| $K_G$ | The maximum reproductive capacity, i.e., reproductive capacity of fully-matured parents of generation $G$ |
| $E_t$ | Epimark level of the parents of a $t$-year-old colony, that is, the time derivative of $C_t$ (Fig. 3B) |
| $\varepsilon$ | Relative strength of epigenetic inheritance |
| $I_t$ | Imprinting level of the offspring produced in year $t$ |
| $A_{i,t}$ | The number of $i$-year-old alates produced in year $t$ |
| $q_{G,i}$ | The reproductive value of the $i$-year-old alate |

**Table S2.** Parameter combinations in figures.

| Figure | Simulation results | Varied<br>Parameters | Fixed<br>Parameters |
| --- | --- | --- | --- |
| Fig. 4A | Evolvability of eusociality | $\beta, F_0, T, \varepsilon$ | |
| Fig. 4B-D | Fitness dynamics | $\beta$ | $F_0, \varepsilon, T$ |
| Fig. 4E | Generations required for the<br>origin of eusociality | $\varepsilon$ | $\beta, F_0, T$ |
| Fig. S7 | Fitness dynamics | $T$ | $\beta, F_0, \varepsilon$ |

